# Aberrant expression of the testis kinase TSSK6activates FAK–STAT3 signaling to promote tumorigenic growth

**DOI:** 10.64898/2026.03.10.710807

**Authors:** Magdalena Delgado, Ian Costello, Sruthi Potturi, Zane Gibbs, Angelique W. Whitehurst

**Affiliations:** Department of Pharmacology, UT Southwestern Medical Center, Dallas, TX 75390, USA

**Keywords:** TSSK6, focal adhesion, colorectal cancer, FAK, STAT3, Cancer testis antigen, PLEKHA7, MEF2C

## Abstract

Cancer–testis antigens (CTAs) are germ cell–restricted proteins aberrantly expressed in diverse malignancies, yet whether these proteins contribute to tumor progression remains poorly defined. The CTA Testis-Specific Serine Kinase 6 (TSSK6) is frequently expressed in colorectal cancer (CRC), where it promotes key hallmarks of tumor progression, including anchorage-independent growth, invasion, and in vivo tumor formation. However, the signaling network by which ectopic TSSK6 drives these phenotypes has not been established. Here, we define the signaling pathways regulated by TSSK6 in CRC cells. We demonstrate that TSSK6 promotes focal adhesion kinase (FAK) activation, leading to enhanced STAT3 Ser727 phosphorylation and increased STAT3-dependent transcription. Transcriptomic analyses revealed induction of gene programs enriched for extracellular matrix remodeling, cytoskeletal organization, adhesion, and motility in a STAT3-dependent manner. Disruption of the FAK–STAT3 signaling axis abrogated anchorage-independent growth and invasive outgrowth in a spheroid model. Collectively, these findings demonstrate that aberrant expression of a germline-restricted kinase promotes focal adhesion signaling, which activates STAT3-dependent transcriptional programs that promote anoikis resistance and invasive behavior in CRC.

## Introduction

Cancer cells frequently anomalously express genes that are normally enriched in germline tissues. These genes, commonly referred to as cancer–testis antigens (CT-antigens, CTAs) or cancer–germline genes, are abnormally expressed across a broad range of malignancies^1, 2^. Several hundred genes have been categorized within this group to date^3–5^. Although their expression in tumors is well documented, the functional contributions of many of these proteins to tumor progression remain incompletely understood.

Recent work from our group and others has demonstrated that aberrantly expressed testis-restricted proteins can be functionally integrated into tumor cell signaling networks to support diverse oncogenic processes. Examples include PRAME, which antagonizes retinoic acid–mediated differentiation signaling^6^; MAGE family proteins, which enhance degradation of tumor suppressor proteins^7^; HORMAD1, which supports radioresistance and mitigates replication stress^8–11^, and FATE1, which alters mitochondrial pro-death signaling^12, 13^. These findings indicate that cancer–testis proteins are not merely bystanders but can actively contribute to malignant progression. Such observations highlight the remarkable plasticity of tumor cells to co-opt proteins with highly tissue-restricted physiological roles to promote neoplastic behaviors.

We recently identified the cancer–testis antigen Testis-Specific Serine Kinase 6 (TSSK6) as aberrantly expressed in approximately 60% of colorectal cancers (CRC)^14^. TSSK6 mRNA expression is associated with reduced overall survival in CRC cohorts. We demonstrated that ectopic expression of TSSK6 promotes anchorage-independent growth, invasion through Matrigel-coated transwell filters, and xenograft tumor formation, whereas depletion of TSSK6 suppresses these phenotypes. Importantly, kinase activity is required for these tumorigenic effects, as expression of a kinase-inactive TSSK6 mutant failed to enhance anchorage-independent growth or invasion^14^.

The TSSK family comprises five members in humans^15^. TSSK6 expression is detected in spermatocytes, spermatids, and mature sperm, with some expression in Sertoli cells^14, 16–18^. Knockout models of three TSSK genes in mice result in severe sperm deformities and male infertility, while mice otherwise appear healthy and develop normally^16–18^. TSSK6 localizes to the head in elongating spermatids and mature sperm, where it has been implicated in actin remodeling^16, 17^. In contrast, in CRC cells we previously observed that TSSK6 localizes to focal adhesions through co-localization with paxillin and tensin at the cell periphery and enhances focal adhesion formation^14^. However, the signaling mechanisms by which TSSK6 regulates focal adhesion dynamics and promotes tumorigenic phenotypes remain unknown.

Here, we define the signaling network downstream of TSSK6 in CRC cells. We demonstrate that TSSK6 promotes activation of focal adhesion kinase (FAK), which is required for downstream STAT3 activation. STAT3 activity is required for expression of genes involved in adhesion, motility, and invasion, including MEF2C and PLEKHA7, which are necessary for TSSK6-driven anchorage-independent growth. In addition to its previously demonstrated effects on transwell invasion, we show that TSSK6 promotes invasive outgrowth of epithelial spheroids into Matrigel–collagen matrix in a FAK-, STAT3-, MEF2C-, and PLEKHA7-dependent manner. Together, these findings reveal that aberrant expression of a sperm-restricted kinase in CRC engages focal adhesion signaling to drive STAT3-dependent transcriptional programs that promote adhesion, motility, and invasion, suggesting a potential role in metastatic progression. Moreover, these data suggest that TSSK6 may represent a previously unrecognized therapeutic vulnerability in CRC.

## Materials and Methods

### Cell Lines and Culture conditions

HCT116 cells were obtained from Cyrus Vaziri (UNC). HT-29, DLD-1, and LOVO cell lines were obtained from Jerry Shay (UT Southwestern). Colorectal cancer cells lines were maintained in high glucose DMEM supplemented with 10% FBS (Cat# F0926, Sigma-Aldrich), 1% antibiotic/antimycotic (Cat# 400-101, GeminiBio) and 2.5 mM HEPES pH7.4 at 37°C in a humidified 5% CO_2_ atmosphere. Human colonic epithelial cells (immortalized with hTERT and CDK4, expressing RASV12 and lacking p53 (HCECs)) were developed by Jerry Shay (UT Southwestern)^19, 20^. TSSK6 overexpressing phenotypes were previously benchmarked^14^. HCECs were maintained in high glucose DMEM supplemented with 5 nM sodium selenite (Cat# S5261, Sigma-Aldrich), 2 ug/mL apo-transferrin (Cat# T1147, Sigma-Aldrich), 20 ng/mL epidermal growth factor (Cat# 236-EG, R&D), 10 ug/mL insulin (Cat# 12585-014, Gibco), 1 ug/mL hydrocortisone (Cat# H0888, Sigma-Alrich), 2% FBS and 1% antibiotic/antimycotic at 37°C in a humidified 5% CO_2_ atmosphere. Stable expressing cell lines were created using lentiviral transduction as previously described^14^. HEK293T cells were obtained from ATCC and cultured in high glucose DMEM supplemented with 10% FBS, 1% anitbiotic/antimycotic and 2.5 mM HEPES pH7.4 at 37°C in a humidified 5% CO_2_ atmosphere. All cells were periodically evaluated for mycoplasma contamination by a mycoplasma PCR detection kit (Cat# G238, ABM). Authenticity of cell lines was evaluated by detection of ten genetic loci using the GenePrint® 10 System (Promega) and cross referencing to ATCC or internal genetic profiles.

### Antibodies and Inhibitors

Antibodies used: TSSK6 (sc-514076, Santa Cruz) (1:500 IB), STAT3 (sc-8019, Santa Cruz (1:1000 IB), STAT3 (9139, CST)(1:1000 IB, 1:300 IF), pSTAT3 (Ser727) (9134, CST)(1:1000 IB), pSTAT3 (Y705) (9131, CST), Vinculin (sc-73614, Santa Cruz) (1:2000 IB), Beta-Tubulin (2128, CST) (1:5000), V5-tag (anti-rabbit,13202, CST) (1:5000 IB), V5-tag (anti-mouse, 46-0705, Invitrogen)(1:2000 IB), PLEKHA7 (PA5-85686, Invitrogen) (1:1000 IB), MEF2C (ab227085, Abcam)(1:5000 IB), pFAK (Y397) (44-624G, Invitrogen) (1:1000 IB, 1:200 IF), FAK (3286, CST)(1:1000 IB). PF-573228, ATP-competitive FAK inhibitor, was purchased from ApexBio Technology (Cat#B1523).

### Transfections

For siRNA transfections, cells were trypsinized and seeded in Opti-MEM containing Lipofectamine™ RNAiMAX (Cat#13778, Thermo Fisher Scientific) complexed with siRNAs at a final concentration of 50 nM for all cells except LOVOs and HT29’s, which were transfected with a final concentration of 100 nM. siRNAs were purchased from Sigma as follows: non-targeting controls (VC30002), TSSK6 pool (SASI_Hs01_00190487, (5’CAGUUGCCCUUGUUCGGAA3’) and SASI_Hs01_00190489 (5’AGACAAACUUCUGAGCG 3’)), STAT3 pool (D-003544-02, D-003544-03, D-003544-04, D-003544-19), PLEKHA7 pool (D-018008-01, D-018008-02, D-018008-03, D-018008-04), MEF2C pool (D-009455-01, D-009455-02, D-009455-03, D-009455-04).

### Plasmids

cDNAs encoding human *TSSK6* were obtained in pDONR223 (DNASU) and cloned into pLX302 or pLX304 (Addgene Plasmid #25896, #25890) using the Gateway Cloning system (Cat#11791 Thermo Fisher Scientific). For control transfections, Hc-Red-V5 in pLX302/pLX304 was used. psPAX2 and pMD2.G lentiviral constructs were purchased from Addgene (Plasmid #12260, #12259).

### Phosphoarray

HCT116 cells were transfected with siCTRL or siTSSK6 as described above for 72 hours. Cells were processed using the Proteome Profiler Human Phospho-kinase Array Kit (Cat#ARY003C, R&D Systems) according to manufacturer’s instructions. Briefly, cells were lysed in provided lysis buffer at a concentration of 1×10^7^ cells/mL, centrifuged, and supernatant collected. Membranes were incubated overnight at 4 degrees Celsius with the supernatant at a concentration of 200 ug/2 mLs. Membranes were washed with buffers, incubated with biotin-conjugated secondary antibodies, and streptavidin-horse radish peroxidase provided with the kit as instructed. Membranes were visualized using chemiluminescence and autoradiography films using a range of exposure times from 1-10 minutes. Positive signal intensity for each spot was quantified using Image J software. Negative control spots were used to subtract background signal from each spot. Average signal was determined for technical duplicate spots representing each phosphorylated kinase protein. Averaged signals were then normalized to positive controls to control for variations in total protein per sample.

### Cell Lysis and Immunoblotting

Samples were lysed in preheated (100°C for five minutes) 2X Laemmli sample buffer with Beta-mercaptoethanol and boiled for six minutes. Samples were resolved using SDS-PAGE and transferred to an Immobilon PVDF membrane (Cat#IPVH00010, Millipore), blocked in 5% non-fat dry milk (total proteins) or 5% BSA (phospo-antibodies) followed by incubation in the respective primary antibodies overnight. For the TSSK6 antibody (sc-514067) membranes were pretreated with SuperSignal^TM^ Western Blot Enhancer (Cat#46640, Thermo Fisher Scientific) and the antibody diluted in buffers provided with the kit. Following incubation, membranes were washed three times with Tris Buffered Saline (20 mM Tris, 150 mM NaCl, 0.1% Tween-20) (TBST) and incubated for either one hour at room temperature or overnight at 4 degrees Celsius with horseradish peroxidase-coupled secondary antibodies (Jackson Immunoresearch). Subsequently, membranes were washed three times with TBST and then developed using SuperSignal™ West Pico PLUS chemiluminescence substrate (Cat#45-000-875, Thermo Fisher Scientific) and imaged using autoradiography films or the Licor Odyssey XF Imaging system. Films were scanned using the EPSON perfection v700 photo scanner.

### Soft agar assays

Cells were suspended into 0.366% bacto-agar and then plated onto solidified 0.5% bacto-agar. Cells were seeded at a density of 500 cells (HCT116) or 3000 cells (DLD-1) per 12-well plate. HCT116 or DLD-1 cells were transfected with siRNA for 48 hours and then collected for plating into soft agar. After 10-14 days, colonies were stained for 1 hour with 0.01% crystal violet in 20% methanol. Background stain was removed by washing 3x in water for 30 minutes and 1x overnight. Images were captured with a Leica S9D dissecting microscope and quantitated by ImageJ software.

### STAT3 Luciferase Reporter Assays

STAT3 Luciferase Reporter Assays were performed in 293T cells using a STAT3 Reporter Kit (Cat#79730, BPS Bioscience) as described by the manufacturer. Briefly, 293T cells were plated in 96-well plates at a density of 15,000 cells per 100 uLs of media. Cells were transfected with 60 ng of STAT3 reporter (STAT3 luciferase reporter + constitutively expressing Renilla luciferase vector), 60 ng of plx302 HcRed or plx302 TSSK6 plasmid, and 0.35 uL Lipofectamine 2000 (Cat#11668027, Thermo Fisher Scientific) in 30 uL of Optimem. After 48 h of incubation, old media was removed, and PBS or 75 ng IL-6/50 uls of media was added to the cells and incubated overnight. Dual-Glo Luciferase Assay (Cat#E2920, Promega) was performed to assess STAT3 transcriptional activity as described by the manufacturer. Firefly luciferase substrate was added to the cells, and firefly luminescence was read using a CLARIOstar Plus plate reader (BMG Labtech). Firefly quench/renilla luciferase substrate was added, and renilla luminescence was read. Firefly luminescence readout was normalized to renilla luminescence to determine STAT3 reporter activity.

### Immunofluorescence Assays

Cells plated on glass coverslips were washed twice with PBS, fixed with 3.7% formaldehyde for 10 minutes at room temperature. Cells were then washed twice with PBS and incubated in 0.25% Triton-100 for 10 minutes prior to washing three times with 1X PBS. Next coverslips were blocked for at least 30 minutes at room temperature or overnight at 4°C in blocking buffer (1% BSA in 0.001%TBST mixed with PBS). Primary antibodies were diluted with blocking buffer and incubated with samples for 1 hour at room temperature. Next, samples were washed three times in blocking buffer for 5 minutes each, followed by incubation with Alexa Fluor-conjugated secondary antibodies (Thermo Fisher Scientific) in blocking buffer at a dilution of 1:1000 for one hour at room temperature. Lastly, samples were washed three times for five minutes with 0.5% BSA in water and 1 time with water, followed by mounting onto glass slides using ProLong™ Gold Antifade reagent with DAPI. Images were acquired by confocal microscope Zeiss LSM700 and processed using Zen 3.8 software (Zeiss). Signal intensity of STAT3 staining or P-FAK foci was quantitated with ImageJ.

### RNA-Sequencing

Total RNA was isolated from HCEC1CTRPs +/- plx302 TSSK6 and HCT116 cells reverse transfected with siCTRL or TSSK6 for 72 h using the GenElute Mammalian Total RNA MiniPrep Kit (Cat#RTN350, Sigma-Aldrich) and treated with DNaseI (Cat#AMPD1, Sigma-Aldrich) as described by the manufacturers. Libraries of mRNA were prepared using the KAPA mRNA Hyper Prep Kit (Cat#KK8580, KAPA Biosystems) as described by the manufacturer. Samples were multiplexed with NEBNext Multiplex Oligos for Illumina adaptors (Cat#E7335, NEB) and 75 bp paired end reads were generated (35-50M per sample) on an Illumina NextSeq 2500 (UTSW McDermott Center Next Generation Sequencing Core). Raw reads were aligned to the human genome assembly (hg38) using Bowtie (ver2.3.2) and the mapped read files were sorted and indexed with SAMtools (ver 1.6). Differential expression analysis was performed using R package DESeq2.

### Enrichr Analysis

Gene lists were generated from HCT116 KD differential expression analysis using log2(fold change) less than –0.5 (p-value < 0.01) and HCEC1CTRP +TSSK6 differential expression analysis using log2(fold) greater than 0.5 (p-value < 0.01). These gene lists were input into Enrichr (https://maayanlab.cloud/Enrichr/) separately for gene set enrichment analysis^21^. Using the Transcription suite of databases, we queried for STAT3 datasets and extracted those with a significant overlap with our TSSK6 knockdown or overexpression gene lists (p-value < 0.05). Using the Pathways and Ontologies suite of databases we surveyed the top ten most enriched datasets from each database and extracted those related to cell migration, invasion, adhesion and the cytoskeleton (p-value < 0.05).

### Gene expression assays

Total RNA was isolated from cells using the GenElute Mammalian Total RNA MiniPrep Kit (Cat#RTN350, Sigma-Aldrich) and treated with DNaseI (Cat#AMPD1, Sigma-Aldrich) as described by the manufacturers. Equal amounts of RNA per experimental condition were reverse transcribed using the Applied Biosystems High-Capacity cDNA Reverse Transfection Kit (Cat#4368814, Thermo Fisher Scientific) with random primers or oligo dT (Cat#12577-011, Thermo Fisher Scientific) as described by the manufacturers. cDNA was used for gene expression analysis performed using Taqman Gene Expression Master Mix (Cat#4369514, Thermo Fisher Scientific) and predesigned Taqman Gene Expression Assays (Thermo Fisher Scientific) on an Applied Biosystems QuantStudio 3 Real-Time PCR instrument. Taqman Gene Expression Assays used are as follows: *TSSK6* (Hs02339769_s1), *CARMIL2* (Hs00418748_m1), *PODXL* (Hs00418748_m1), *PLEKHA7* (Hs00957677) *MEF2C* (Hs00231149_m1) *STAT3* (Hs00374280_m1), *RPL27* (Hs03044961_g1).

### 2.5D Spheroid Growth Assays

Cells were pre-plated into 10 cm dishes so they were 90-100% confluent when plating into spheroid growth culture. For RNAi experiments, cells were reverse transfected with siRNA for 48 h and then plated into spheroid growth cultures. Rat tail collagen I (Cat#CB-40236, Thermo Fisher Scientific) was activated by mixing 500 uL collagen with 62.5 uL 10X PBS, 62.5 uL 0.1 N NaOH, and 10 uL of 1:100 HCl. Activated collagen was then mixed 1:1 with Growth Receptor Free Matrigel (Cat#354230, Thermo Fisher Scientific) while keeping on ice. Matrigel/collagen was plated into 96-well plates at 50 uL per well pipetting slowly to spread evenly and thinly without introducing bubbles. Plates were incubated in humidified growth incubators for at least 30 min to allow for solidification of the Matrigel/collagen. Cells were trypsinized, rinsed with PBS and resuspended in growth media. Cells were counted and resuspended in 2% matrigel assay media with 5 ng/mL of EGF at a concentration of 2500 cells/ 200 uL media for DLD-1s and 1250 cells/200 uL media for HCEC1CTRP for each well. For FAK inhibitor experiments, PF-573228 (500 nM) or control DMSO (0.005%) was added to the 2% matrigel assay media prior to cell resuspension. Cells were allowed to incubate for 120 hours and then fixed with 2% formaldehyde in PBS for at least 1 hour at room temperature. Spheroids were imaged using an Olympus IX51 at 20X on brightfield, and circularity was measured using ImageJ.

### Statistics

Data are presented as mean ± SD of independent biological replicates as indicated in figure legends. Technical replicates (e.g., multiple wells or repeated measurements within the same experiment) were averaged prior to statistical analysis. For assays in which conditions were performed in parallel within the same experiment, paired analyses were used. When data were approximately normally distributed, parametric tests (paired or one-sample t-tests, as appropriate) were applied. When distributions deviated from normality as judged by d’Agostino test the corresponding nonparametric tests were used (Wilcoxon signed-rank test for paired comparisons; Mann–Whitney test for unpaired comparisons). Circularity and colony-level measurements were analyzed at the level of independent biological experiments; individual colony measurements are shown to illustrate distribution but were not treated as independent replicates for statistical testing. All statistics are calculated for paired, one sample t-test, two-tailed unless otherwise specified. Statistical analyses were performed using GraphPad Prism.

## Results

To define signaling pathways regulated by TSSK6 in CRC cells, we performed a loss-of-function phospho-signaling screen (Figure 1A). We assessed changes in the phosphorylation state of 37 signaling proteins, as well as total levels of HSP60 and β-catenin, following TSSK6 depletion. This analysis was conducted in HCT116 colorectal cancer cells, which express endogenous TSSK6 and require its expression for anchorage-independent growth.

**Figure 1.**
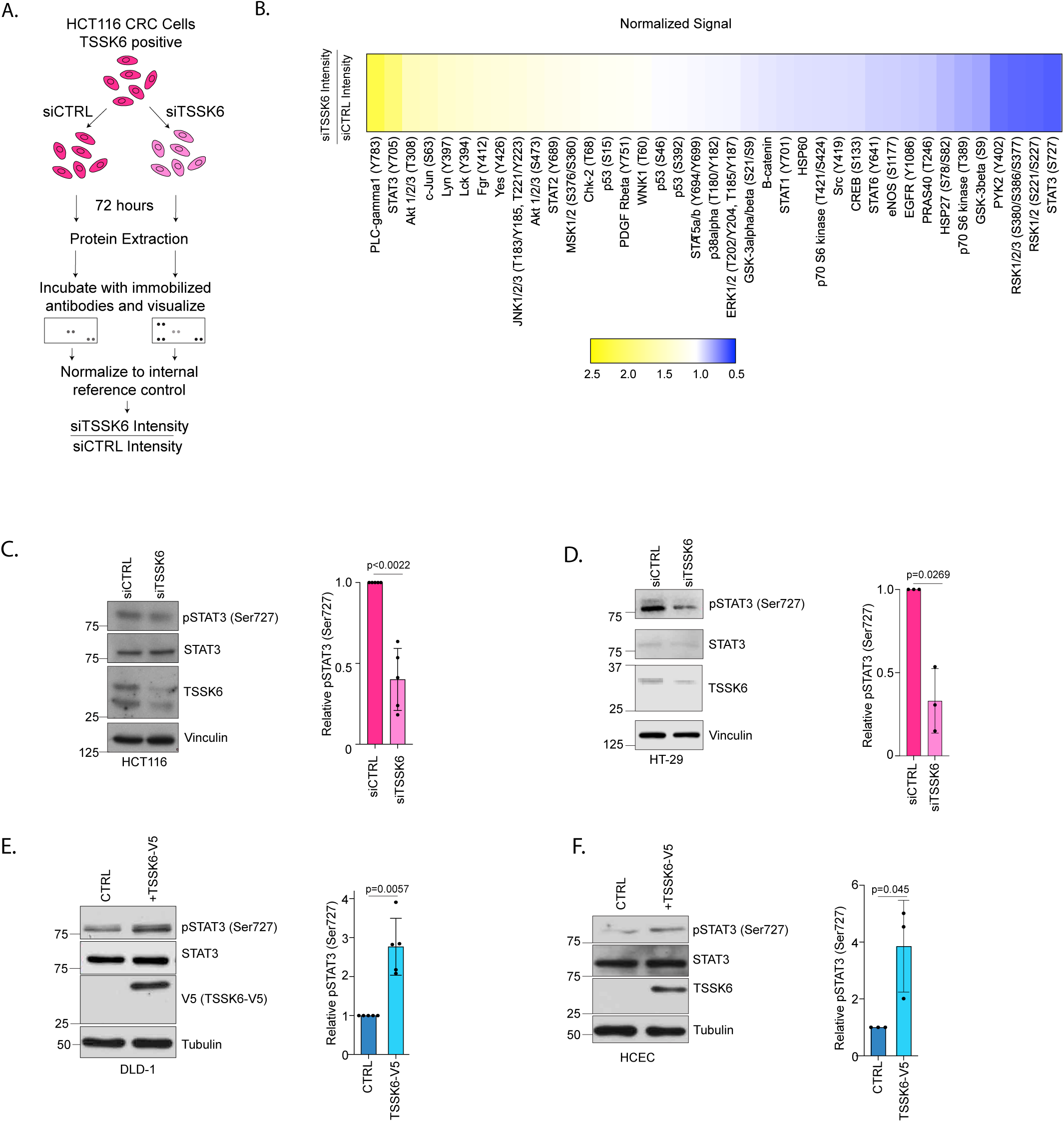
TSSK6 supports multiple signaling nodes. A. HCT116 were transfected with siCTRL or siTSSK6 for 72 hours. Lysates were evaluated for phosphorylation using phosphoarray membranes. Values are normalized to siCTRL, n=2, each with technical replicates. B. Heat map representing the average normalized signal across both replicates. C. HCT116 (n=5) were transfected with siTSSK6 for 72 hours, lysed, resolved by SDS-PAGE and immunoblotted with indicated antibodies. HT-29 (n=3) were transfected with siTSSK6 for 96 h, lysed, resolved by SDS-PAGE and immunoblotted with indicated antibodies. E. DLD-1s ± TSSK6-V5 (n=5) were lysed, resolved by SDS-PAGE and immunoblotted with indicated antibodies. F. As in E for HCECs ± TSSK5-V5 (n=3).

Depletion of TSSK6 resulted in a ≥40% reduction in phosphorylation of STAT3 (Ser727), RSK1/2/3 (multiple sites), and PYK2 (Tyr402), returning these kinases among the most strongly altered signaling nodes in the phospho-screen (Figure 1B; Supplemental Table 1). Immunoblot validation confirmed reduced phosphorylation of each of these proteins upon TSSK6 depletion in HCT116 cells (Figure 1C, Supplemental Figure 1A,B). Given the established role of STAT3 in anchorage-independent growth and tumor progression, we prioritized STAT3 for further mechanistic investigation^22–26^.

Consistent with the phosphoarray results, depletion of TSSK6 in HCT116 cells led to a marked reduction in STAT3 Ser727 phosphorylation, as assessed by immunoblotting (Figure 1C). A similar decrease was observed in HT-29 cells, which also express endogenous TSSK6 (Figure 1D), indicating that regulation of STAT3 by TSSK6 occurs across multiple colorectal cancer cell lines.

To determine whether TSSK6 is sufficient to induce STAT3 phosphorylation, we extended our analysis to gain-of-function systems. Introduction of TSSK6 into DLD-1 cells, which lack endogenous TSSK6, resulted in a 2–3-fold increase in STAT3 Ser727 phosphorylation (Figure 1E). This phenotype was also recapitulated in immortalized human colonic epithelial cells (HCEC), which similarly lack endogenous TSSK6 (Figure 1F).

In addition to changes in STAT3 Ser727 phosphorylation, the phosphoarray analysis suggested altered phosphorylation of STAT3 at Tyr705 following TSSK6 depletion. This phenotype was recapitulated in independent transfections in HCT116 cells (Supplemental Figure 1C), but was highly variable in in additional cell lines (data not shown) suggesting that regulation of individual STAT3 phosphorylation sites downstream of TSSK6 may be influenced by cellular context. Despite this variability, modulation of STAT3 Ser727 was a consistent feature of both loss- and gain-of-function models, leading us to focus subsequent analyses on STAT3 as a downstream effector of TSSK6.

STAT3 is a transcription factor frequently activated in a wide array of malignancies, where it is required for diverse oncogenic behaviors ^23–29^. Indeed, depletion of STAT3 from HCT116 cells led to a dramatic decrease in soft agar colony-forming ability (Figure 2A). To determine whether STAT3 is required for TSSK6-induced anchorage-independent growth, we depleted STAT3 in DLD-1 cells overexpressing TSSK6 and evaluated soft agar growth. TSSK6-enhanced soft agar growth was reversed in the absence of STAT3 (Figure 2B). Because STAT3 functions primarily as a transcription factor, we next asked whether TSSK6 expression was sufficient to enhance STAT3-dependent transcriptional activity. Using a luciferase-based reporter driven by a canonical STAT3 response element, we found that expression of TSSK6 increased reporter activity approximately four-fold under basal conditions and further enhanced reporter activity (∼2-fold) following JAK-dependent STAT3 activation (Figure 2C). To further assess STAT3 activation status, we examined STAT3 subcellular localization. Stable expression of TSSK6 in DLD-1 cells was associated with increased nuclear accumulation of STAT3, consistent with enhanced STAT3 activation (Figure 2D). Together, these data indicate that STAT3 activity is required for TSSK6-driven anchorage-independent growth and that TSSK6 expression is sufficient to enhance STAT3 functional output.

**Figure 2.**
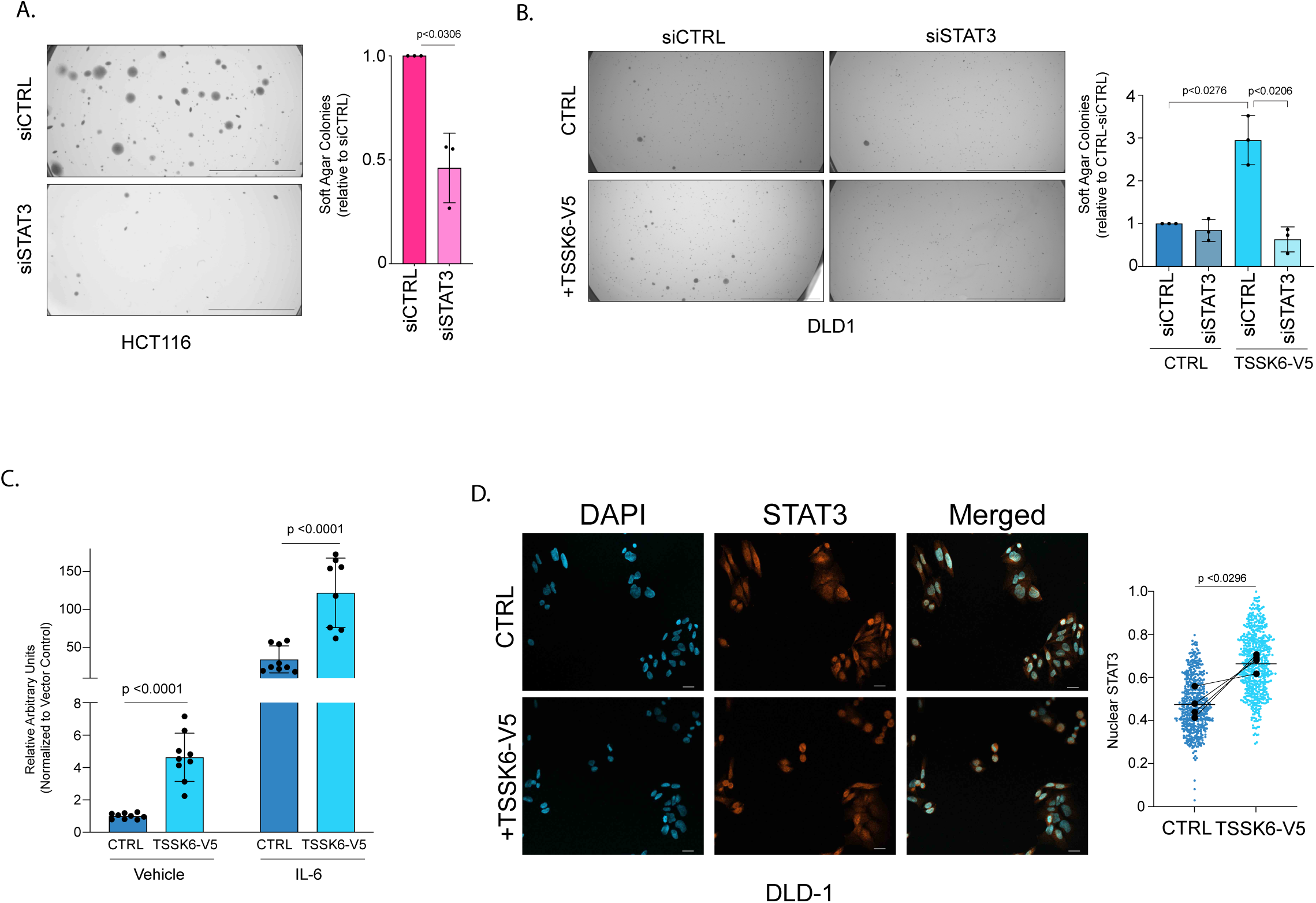
STAT3 is required for TSSK6-induced phenotypes. A. HCT116 (n= 10) or B. DLD-1 ± TSSK6-V5 (n=9) were transfected with siCTRL or siSTAT3 for 48 hours then plated into soft agar. 10-14 days later, samples were stained and quantitated. Bars represent mean ± s.d. C. 293Ts were transfected with STAT3 reporter ± TSSK6-V5 (n=9). After 48 hours, vehicle control or 75 ng/uL Il-6 was added, and cells were incubated overnight. Dual Glo Luciferase assays were used to read STAT3 reporter firefly luminescence normalized to constitutively active renilla luminescence. Samples with IL-6 were analyzed with a paired t-test. D. Confocal images of DLD-1 ± TSSK6-V5 cells plated onto glass coverslips for 1 day prior to staining with STAT3. The scale bar is 20 µm. Nuclear STAT3 signal normalized to total STAT3 signal per cell was quantified using ImageJ. Data points represent individual cells from 3 fields of view across 4 coverslips. Paired t-test used for statistical analysis.

To broadly define transcriptional programs modulated by TSSK6 in CRC, we examined transcriptomes following depletion of TSSK6 in HCT116 CRC cells and stable expression of TSSK6 in non-transformed human colonic epithelial cells (HCEC) (Figure 3A). Volcano plots from each dataset demonstrated widespread and statistically significant transcriptional changes following manipulation of TSSK6 expression (Supplemental Figure 2A). Enrichr analysis identified multiple gene sets involved in extracellular matrix remodeling, cell migration and adhesion, and EMT (Supplemental Figure 2B, Supplemental Table 2). Intersection of the two datasets identified six genes that underwent reciprocal expression changes (log₂ fold change > 0.5 in HCEC and < −0.5 in HCT116; p-value < 0.01) (Figure 3B). All six genes have been implicated in invasion, migration, modulation of cell adhesion and/or actin dynamics (CARMIL2, PODXL, PLEKHA7, MEF2C, COL4A6, PLPPR4^30–41^).

**Figure 3.**
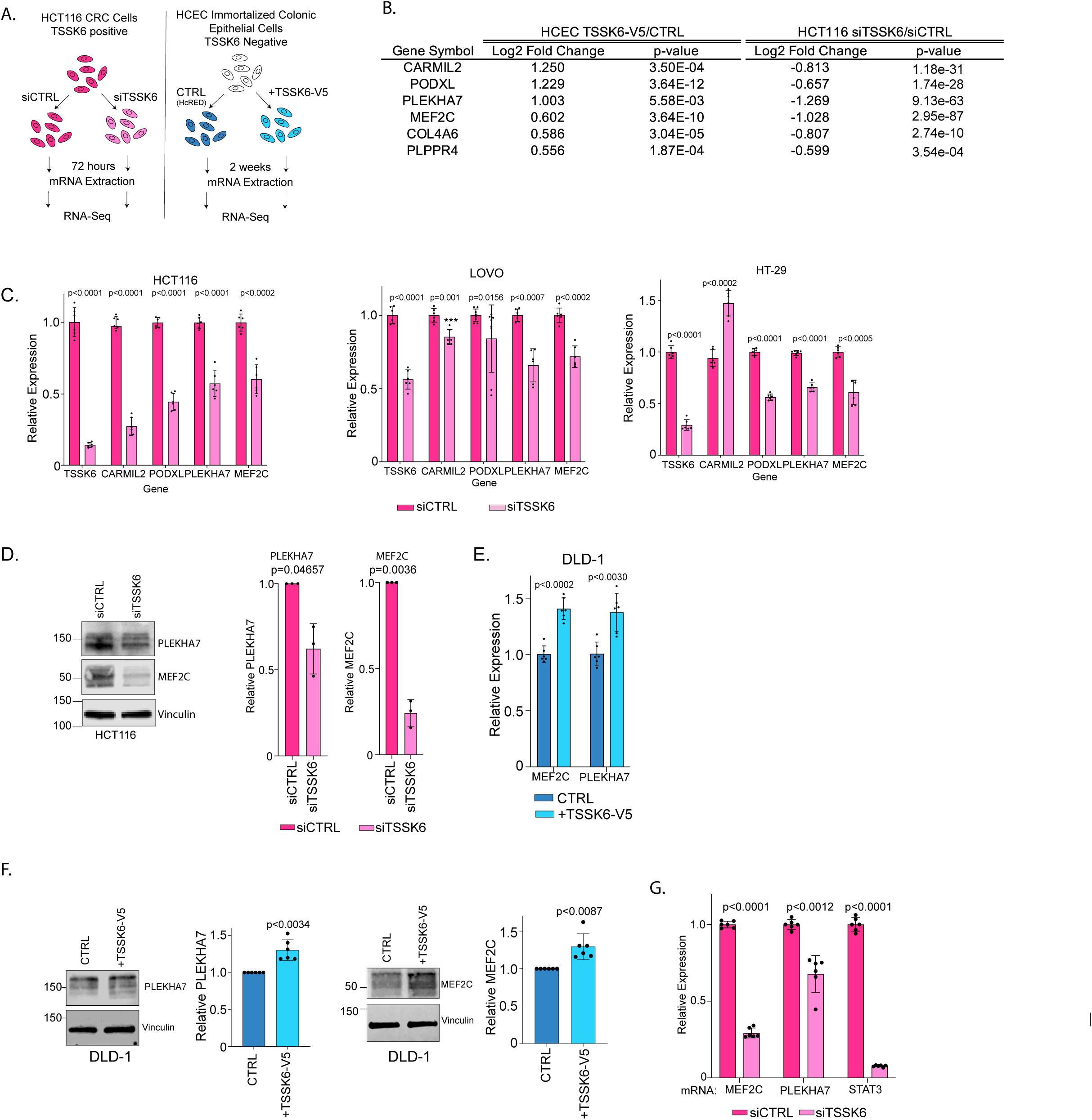
TSSK6 promotes expression of genes involved in ECM, motility and invasion. A. HCT116 were transfected with siCTRL or siTSSK6 for 72 hours and HCECs were stably transfected with plx302 HcRed or plx302 TSSK6-V5. Total RNA was extracted; mRNA libraries were made and multiplexed with Illumina adaptors. RNA-sequencing and differential expression analysis were performed across three replicates for each experimental condition. B. Table of genes that overlapped across the gain-of-function (HCEC, log2(fold change)>0.5 and p-value<0.01) and loss-of-function (HCT116, log2(fold change) <-0.5 and p-value<0.01) differential gene expression analysis. C. HCT116 (n=6) was transfected with siCTRL or siTSSK6 for 72 hours while HT29 (n=6) and LOVO (n≥6) were transfected with siCTRL or siTSSK6 for 96 hours. Cells were lysed, RNA isolated, cDNA generated, and RT-qPCR was performed for the indicated genes. Statistical analysis by one sample t-test, except for PODXL in LOVO’s, where a Wilcoxon ranked test was used due to non-normal distribution. D. HCT116 (n=3) were transfected with siTSSK6 for 72 hours, lysed, resolved by SDS-PAGE and immunoblotted with indicated antibodies. E. DLD-1s ± TSSK6-V5 (n=6) were lysed, RNA isolated, cDNA generated, and RT-qPCR was performed for the indicated genes. F. DLD-1s ± TSSK6-V5 (n=6) were lysed, resolved by SDS-PAGE and immunoblotted with indicated antibodies. G. HCT116 (n=6) were transfected with siCTRL or siSTAT3 for 96 hours and lysed. RNA was isolated, cDNA generated, and RT-qPCR was performed for the indicated genes.

We further evaluated the four genes that exhibited the most substantial expression changes (CARMIL2, PODXL, PLEKHA7, and MEF2C) across three CRC cell lines that endogenously express TSSK6 (HCT116, HT-29, and LOVO)^14^. Among these candidates, PLEKHA7 and MEF2C mRNA levels were consistently dependent on TSSK6 in all three cell lines evaluated (Figure 3C). These transcriptional changes were accompanied by reduced protein accumulation in HCT116 cells depleted of TSSK6 (Figure 3D). Notably, depletion of either PLEKHA7 or MEF2C reduced anchorage-independent growth in HCT116 cells (Supplemental Figure 2D). We next assessed PLEKHA7 and MEF2C expression in a gain-of-function context using DLD-1 CRC cells, which lack endogenous TSSK6. Ectopic expression of TSSK6 resulted in a statistically significant increase in both mRNA and protein levels of PLEKHA7 and MEF2C (Figure 3E,F). Given that STAT3 functions downstream of TSSK6, we next examined whether STAT3-associated transcriptional signatures were enriched in the RNA-seq datasets using focused Enrichr analysis. In both loss- and gain-of-function datasets, multiple STAT3-associated gene sets were significantly enriched (Supplemental Figure 2C). We therefore tested whether PLEKHA7 and MEF2C are modulated by STAT3. Indeed, depletion of STAT3 in HCT116 cells resulted in reduced mRNA accumulation of both genes (Figure 3G). Together, these analyses support a model in which TSSK6 promotes a STAT3-associated transcriptional program.

We previously reported that ectopically expressed TSSK6 localizes to focal adhesions in colorectal cancer cells^14^. To determine whether TSSK6 localization is associated with activation of focal adhesion signaling, we examined focal adhesion kinase (FAK) phosphorylation in both loss- and gain-of-function settings. Ectopic expression of TSSK6 resulted in increased accumulation of phosphorylated FAK, whereas depletion of TSSK6 exhibited a trend in decreasing pFAK (Figure 4A,B). Consistent with enhanced focal adhesion signaling, expression of TSSK6 accelerated cell spreading in DLD-1 cells and increased the proportion of cells exhibiting p-FAK–positive focal adhesion structures at the cell periphery (Figure 4C). FAK activation has previously been linked to STAT3 phosphorylation at both the Ser727 and Tyr705 residues^42, 43^. We therefore examined whether FAK activity contributes to STAT3 phosphorylation downstream of TSSK6. Pharmacologic inhibition of FAK reduced STAT3 Ser727 phosphorylation in HCT116 cells (Figure 4D). Importantly, TSSK6-induced STAT3 Ser727 phosphorylation in DLD-1 cells was abrogated upon FAK inhibition (Figure 4E). Together, these findings indicate that ectopic expression of TSSK6 in CRC cells promotes FAK activation, which is required for downstream STAT3 Ser727 phosphorylation.

**Figure 4.**
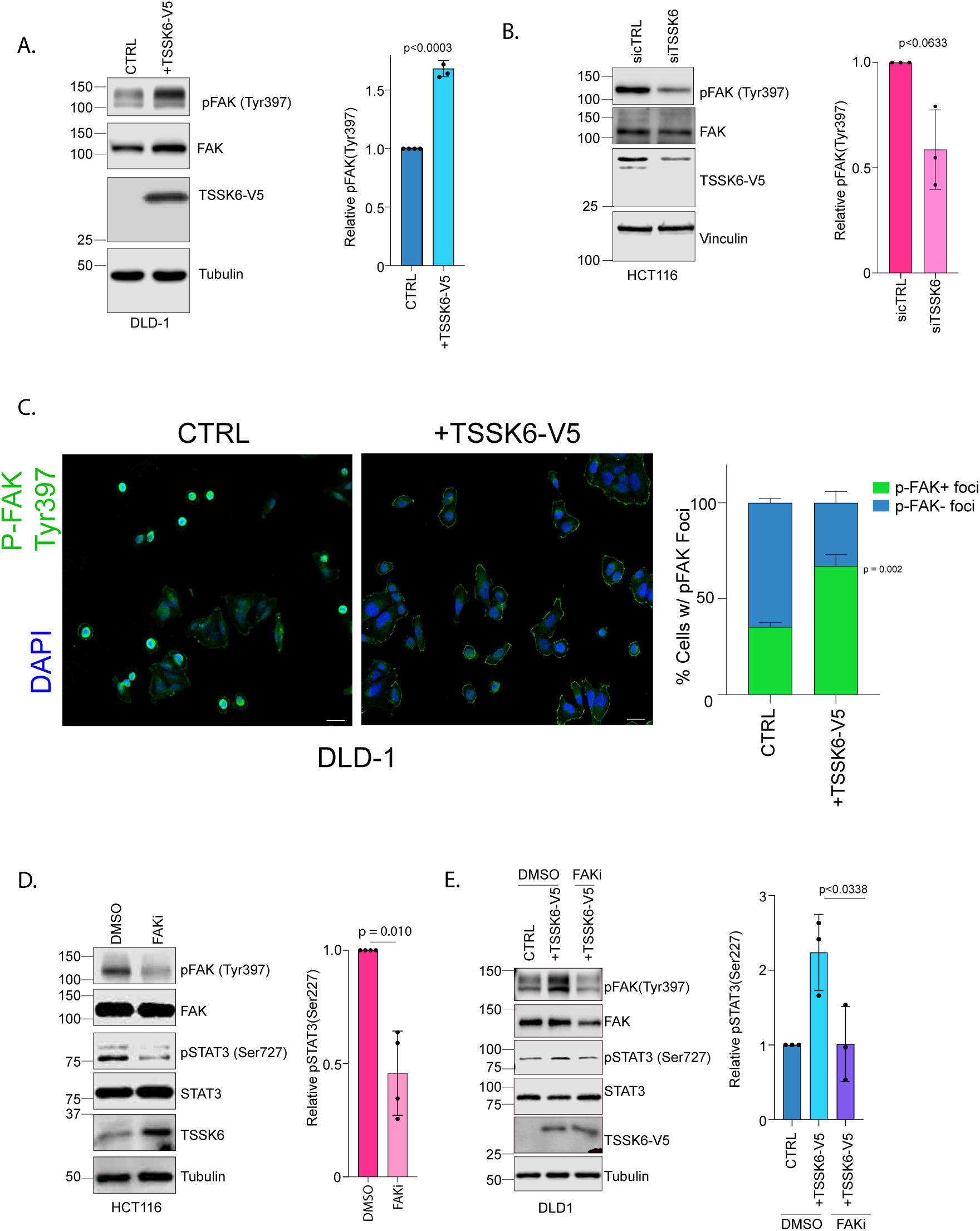
STAT3 activation is dependent on FAK activation. A. HCT116 (n=3) were transfected with siTSSK6 for 72 hours, lysed, resolved by SDS-PAGE and immunoblotted with indicated antibodies. B. DLD-1s ± TSSK6-V5 (n=3) were plated and then lysed after 6 hours, resolved by SDS-PAGE and immunoblotted with indicated antibodies. C. Confocal images of DLD-1 ± TSSK6-V5 cells plated onto glass coverslips for 6 hours prior to staining with P-FAK. The scale bar is 20 µm. The percentage of cells per field of view that were either positive or negative for p-FAK foci were manually quantitated. Data represent cells from 3 fields of view across 4 coverslips. D. HCT116 (n=4) or E. DLD-1 (n=3) were treated with 500 nM PF-573228 for 1 hour, lysed, resolved by SDS-PAGE and immunoblotted with indicated antibodies. Paired t-test was used to analyze +TSSK6-V5 samples in E.

The phenotypes observed following aberrant expression of TSSK6 implicate this kinase in promoting cell adhesion and motility in cancer cells, suggesting a potential role in metastatic behavior. To directly test this possibility, we employed a 2.5-dimensional spheroid invasion assay that assesses invasive outgrowth into a surrounding Matrigel–collagen matrix (Figure 5A). Expression of TSSK6 in cells that lack endogenous TSSK6, including immortalized human colonic epithelial cells and DLD-1 CRC cells, resulted in increased extrusion of cells into the surrounding matrix, as evidenced by loss of spheroid circularity and increased invasive outgrowth (Figure 5B,C). FAK activity was required for this process, as pharmacologic inhibition of FAK nearly completely reversed the TSSK6-induced extrusion phenotype (Figure 5D). STAT3 was also essential for TSSK6-driven invasion, as depletion of STAT3 similarly abrogated spheroid extrusion into the matrix (Figure 5E). Finally, depletion of either MEF2C or PLEKHA7 was sufficient to partially reverse the invasive phenotype induced by TSSK6 expression (Figure 5F). Together, these findings indicate that aberrant expression of TSSK6 promotes a motile and invasive phenotype in cancer cells through activation of focal adhesion signaling, leading to STAT3 activation and induction of transcriptional programs that support invasive behavior.

**Figure 5.**
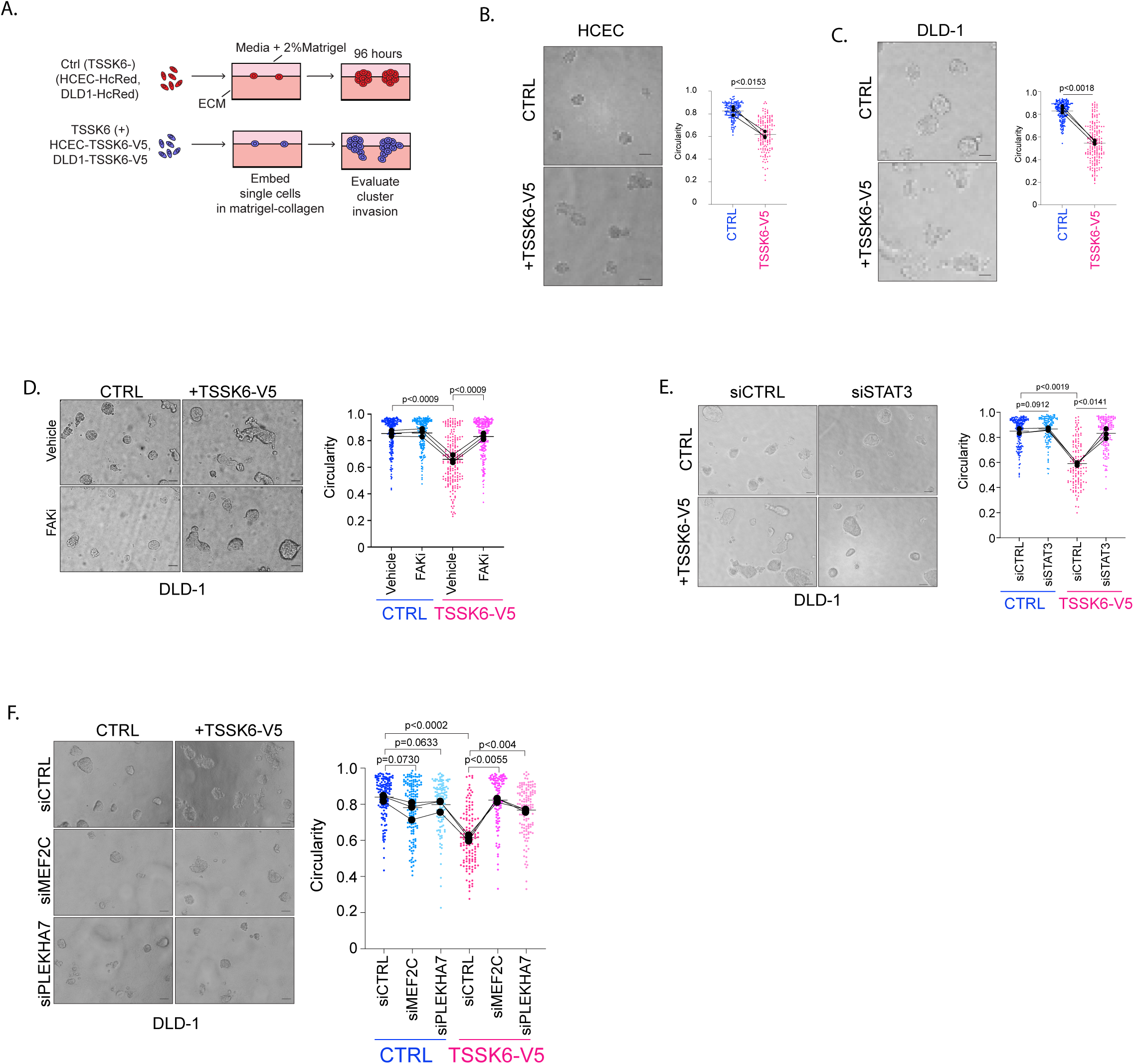
TSSK6 promotes invasive outgrowth. A. DLD-1 or HCECs ± TSSK6-V5 were placed into single cell suspension, embedded into matrigel–collagen, overlayed with 2% matrigel media, and allowed to grow for 120 hours. Cell clusters were evaluated for invasive growth. B and C. Data points represent individual cells from 3-5 fields of view from 2-3 individual wells across 3 plates. D. DLD-1 ± TSSK6-V5 were embedded into matrigel-collagen, overlayed with 2% matrigel media containing either DMSO control or FAK inhibitor and allowed to grow for 120 hours. Data points represent individual cells from 3-5 fields of view from 2-3 individual wells across 3 plates. E and F. DLD-1 ± TSSK6-V5 were transfected with indicated siRNA for 48 h, embedded into matrigel-collagen and allowed to grow for 120 hours. Data points represent individual cells from 3-5 fields of view from 2-3 individual wells across 3 plates.

## Discussion

Sperm undergo extraordinary morphological changes during their development from primordial germ cells to mature spermatozoa. This process requires tissue-specific proteins that mediate dynamic cytoskeletal remodeling necessary for elongation, nuclear condensation, and formation of a motile flagellum^16, 18, 44^. TSSK6 is essential for these morphological transitions; mice lacking TSSK6 exhibit altered nuclear morphology, bent tails, and severely compromised head–tail attachment^16, 18, 44^. These deformities are due to changes in cytoskeletal proteins including actin, tubulin and the sperm structures comprised of these components^16, 18, 45^. Similar phenotypes are observed following loss of the single TSSK family member in zebrafish and Drosophila, suggesting conservation of function across species^45, 46^. Thus, TSSK6 and related family members are essential regulators of cytoskeletal organization during spermatogenesis.

Cancer cells likewise exhibit extensive cytoskeletal dysregulation to accommodate changes in adhesion, motility, and survival following substrate detachment. A defining hallmark of malignant progression is the capacity to bypass anoikis and survive in the absence of attachment^47^. Our findings here and in prior work suggest that TSSK6 expression is commandeered by cancer cells to promote anchorage-independent growth and invasion. We now elaborate this mechanism by demonstrating that TSSK6 enhances focal adhesion kinase (FAK) activation, leading to increased STAT3 transcriptional activity and induction of gene programs enriched for extracellular matrix organization, cytoskeletal remodeling, adhesion, and motility. Thus, TSSK6 appears to integrate into and induce a canonical FAK–STAT3 signaling axis that is frequently aberrantly activated in cancer cells.

Our studies indicate that TSSK6 preferentially enhances phosphorylation of STAT3 at Ser727. Amplified Ser727 phosphorylation has recently been associated with activation of gene programs linked to adhesion, proliferation, and migration in cancer^22^. STAT3 Ser727 phosphorylation represents a convergence point for multiple kinase pathways linked to adhesion and growth factor signaling. Notably, our phospho-signaling screen revealed that TSSK6 depletion reduces phosphorylation of kinases within these pathways, including Src (Y416), EGFR (Y1086), and mTORC1-associated substrates such as p70S6K (T389). The modest yet coordinated reduction in phosphorylation across these nodes suggests that TSSK6 supports a broader adhesion-linked signaling module rather than a single linear cascade. This signaling profile raises the possibility that multiple kinases may contribute to STAT3 Ser727 phosphorylation downstream of TSSK6. However, because FAK inhibition reverses TSSK6-associated STAT3 Ser727 phosphorylation, our data place FAK upstream of STAT3 in this regulatory axis.

The mechanism by which TSSK6 promotes FAK activation remains unclear. In prior work, we demonstrated that kinase-inactive TSSK6 mutants fail to induce anchorage independence, indicating that catalytic activity is required for its tumorigenic function. In germ cells, TSSK6 has been associated with phosphorylation of cytoskeletal regulators, and its loss alters phosphorylation of proteins involved in axonemal organization^18^. Direct substrates of TSSK6 are unknown in sperm and cancer cells. As substrates emerge for TSSK6, it will be possible to delineate whether these events contribute to FAK activation or instead reflect downstream consequences of altered adhesion signaling. Given that TSSK6 localizes to focal adhesions, this activity may occur locally to influence integrin-dependent signaling^14^. The identification of downstream signaling outputs described here provides a framework for future studies aimed at defining direct TSSK6 substrates in cancer cells.

Collectively, these findings highlight the capacity of tumor cells to engage primordial germline programs to acquire neomorphic phenotypes. Aberrant expression of a testis-restricted kinase such as TSSK6 may facilitate escape from anchorage-dependent growth constraints and promote invasive behavior, potentially enhancing early steps of metastatic dissemination. Given the restricted expression of TSSK6 to testis and tumors, targeting this kinase could represent a selective therapeutic strategy with a favorable therapeutic window.

## Author contributions

A.W.W., M.D., conceptualization; A.W.W., M.D., methodology; A.W.W.,I.C., Z.G.,S.P.,M.D., investigation; A.W.W. and M.D. writing—original draft, review & editing; A.W.W., supervision; A.W.W. funding acquisition.

## Supplemental Figure Legends

**Supplemental Figure 1.**
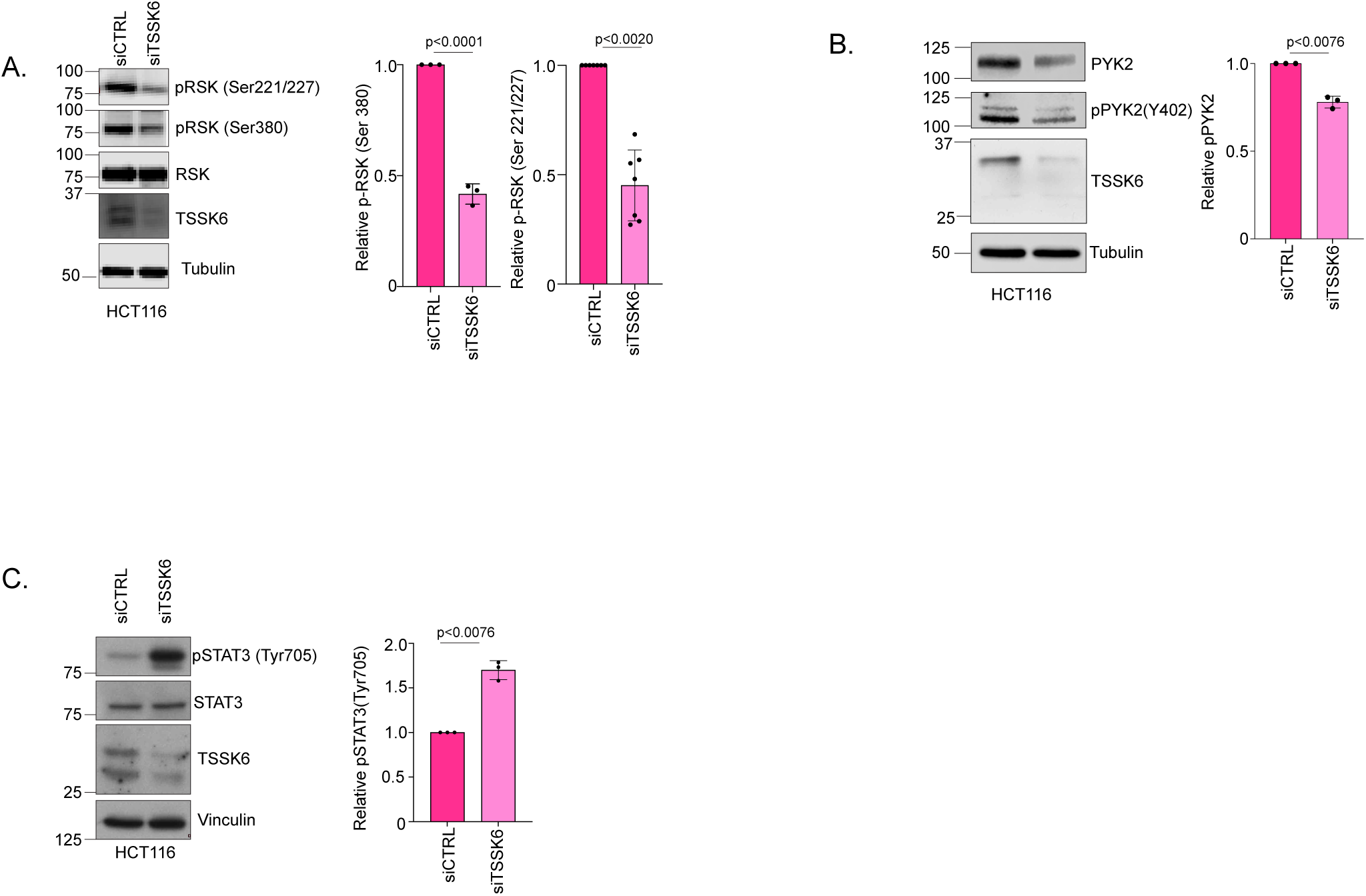
TSSK6 modulates phosphorylation of RSK, PYK2, and STAT3. A, B, C. HCT116 (n=3) were transfected with siTSSK6 for 72 hours, lysed, resolved by SDS-PAGE and immunoblotted with indicated antibodies.

**Supplemental Figure 2.**
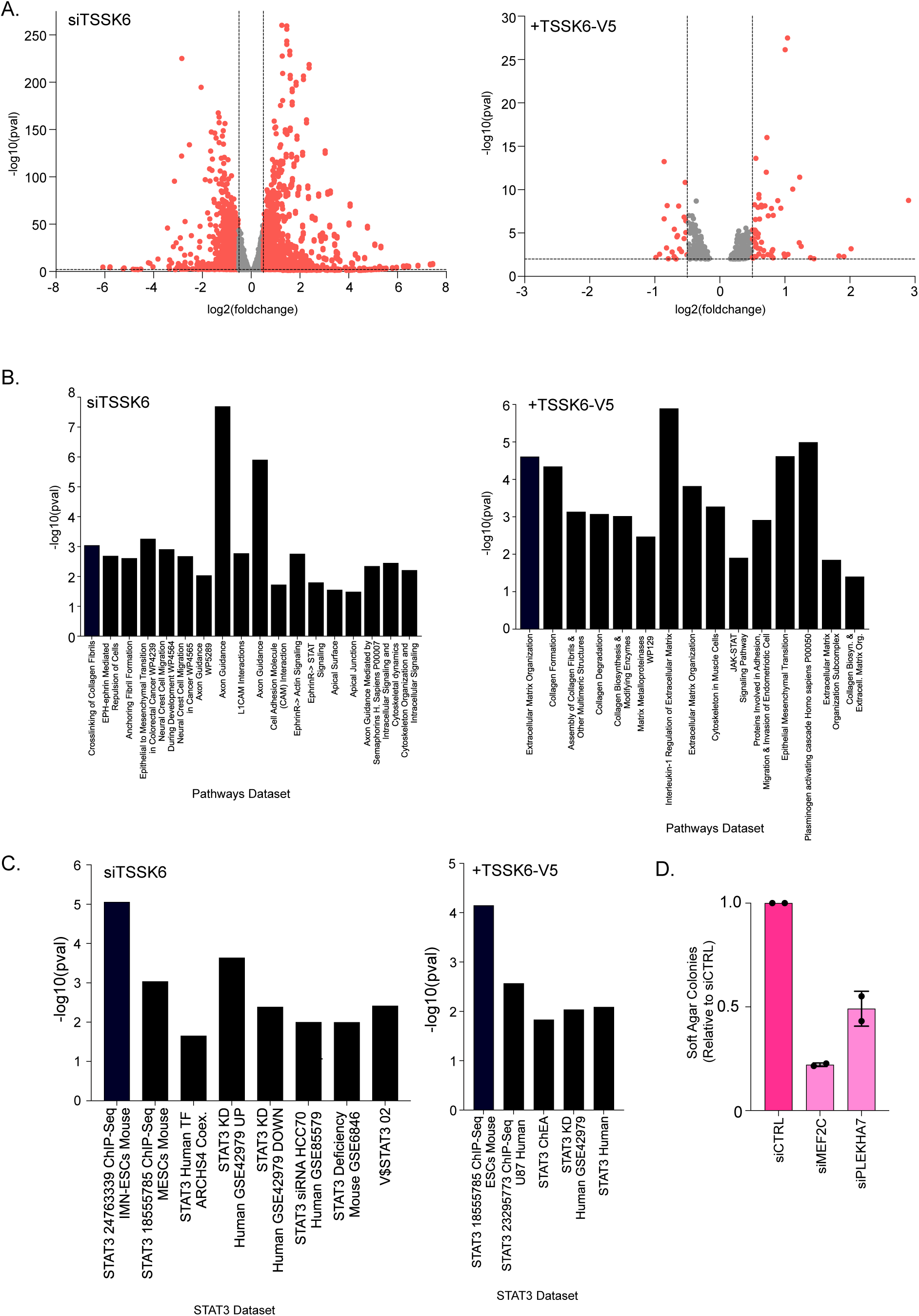
TSSK6 modulates broad transcriptional programs. A. Volcano plots from differential gene expression analysis from loss-of-function (HCT116) and gain-of-function (HCEC) RNA-sequencing. Vertical dotted lines indicate log2(fold change) equals -0.5 or 0.5 and horizontal dotted lines indicate –log10(p-value) equals 2. Data points below this horizontal dotted line are not shown. B. Datasets returned in the Top Ten of Enrichr’s Pathways suite of databases as being enriched for in gain-of-function (HCEC, log2(fold change)>0.5 and p-value<0.01) and loss-of-function (HCT116, log2(fold change) <-0.5 and p-value<0.01) differential gene expression analysis. C. STAT3 datasests returned from Enrichr Transcription suite of databases as being enriched for in gain-of-function (HCEC, log2(fold change)>0.5 and p-value<0.01) and loss-of-function (HCT116, log2(fold change) <-0.5 and p-value<0.01) differential gene expression analysis. D. HCT116 were transfected with indicated siRNA for 48 hours then plated into soft agar. 10-14 days later, samples were stained and quantitated.

## Supplemental Tables

Supplemental Table 1. Phospho-array data from two individual experiments. The mean signal intensity from two technical replicates for each phospho-site is listed for each individual experiment, with the average of both, and average deviation.

Supplemental Table 2. Enrichr analysis from both gain-of-function and loss-of-function RNA-sequencing data. Sheet 1 is the gene list from loss-of-function (HCT116, log2(fold change) <-0.5 and p-value<0.01) differential gene expression analysis. Sheet 2 is the gene list from gain-of-function (HCEC, log2(fold change)>0.5 and p-value<0.01). Sheet 3 is the Enrichr datasets returned from the loss-of-function dataset and Sheet 4 are datasets returned from the gain-of-function dataset.

